# Focused Ultrasound Neuromodulation of the Central Lateral Thalamus in a Non-Human Primate Model of Disorders of Consciousness

**DOI:** 10.64898/2026.09.15.751608

**Authors:** Carter Lybbert, Caroline Garrett, Taylor Webb, Keisuke Tsunoda, Ruksana Begum, Jan Kubanek

## Abstract

**Objectives:** Disorders of consciousness (DOC) affect an estimated 200,000 patients in the United States. Effective treatment options have remained elusive. Focused ultrasound neuromodulation (FUS) of brain regions controlling consciousness, such as the central lateral thalamus (CLT), has emerged as a promising non-invasive treatment. However, prior studies have been limited by broad effective stimulation volumes, missing control conditions, and limited target validation. This study investigated the effects of FUS of the CLT in a nonhuman primate model of DOC with robust control conditions and more robust targeting than previous experiments.

**Materials and Methods:** Two male rhesus macaques underwent 1-hour propofol anesthesia sessions as a model of DOC. Bilateral CLT targets were sonicated using a stereotactically mounted 256-element phased-array focused ultrasound system. Focused and unfocused (random phases) paradigms were tested in both animals, and a spotlighting (sonicating multiple targets) paradigm was tested in one animal. Arousal was monitored via EEG beta power, pulse oximetry, breathing rate, and movement, and compared between sonication and control sessions for each paradigm.

**Results:** Paired Wilcoxon signed-rank tests showed no significant differences for any of the four measures across the three paradigms (all **p**_**FDR**_ ≥ **0.30**). The largest observed effect was increased body movement under the unfocused paradigm (*r*_rb_ = +0.58), though this did not reach statistical significance. Animals remained physiologically stable throughout sessions and recovery periods, with no clinically significant changes in heart rate, breathing rate, blood oxygen saturation, or other safety concerns.

**Conclusions:** Under the tested parameters, FUS of the CLT did not produce measurable changes in arousal. The absence of adverse physiological effects supports the safety of this FUS paradigm. The largest effect occurring under spatially distributed sonication, consistent with prior evidence that broader stimulation volumes produce stronger arousal responses, suggests that distributed network stimulation may warrant investigation as an alternative to focal targeting. Future studies should investigate this possibility and should include robust control conditions to properly quantify effects.

## Introduction

An estimated 200,000 patients are living with prolonged disorders of consciousness (DOC) in the United States.^1^ The lifetime cost of treating a DOC can exceed $1 million, ^1,2^ and puts a significant financial and emotional burden on the patients’ family and caregivers.^3^ DOC is characterized by a persistent, significant impairment of arousal and/or awareness and usually develops after events like stroke, traumatic brain injury, or cardiac arrest.^4^ DOC includes conditions like coma, which involves no arousal or awareness, vegetative state, which presents with wakefulness without awareness, and minimally conscious state, which shows inconsistent awareness.^4^

Consciousness is a complex phenomenon that arises from interactions between multiple neural networks throughout the brain, including thalamocortical and frontoparietal networks, the ascending reticular activating system (involved in arousal), and cortical regions like the default mode network (involved in awareness).^4^ According to the mesocircuit model, the central lateral thalamus (CLT) together with the striatum and prefrontal cortex create a crucial loop which facilitates recovery of consciousness by reactivating these pathways after loss of consciousness.^5^ In this way, the CLT is a key region in the networks controlling recovery of consciousness.

Propofol anesthesia bears many similarities to natural loss of consciousness, including widespread suppression of high-frequency neural activity and disruption of brain-wide network interactions. These changes are reflected by suppressed high-frequency EEG activity, and heightened low-frequency EEG power alongside diminished frontal alpha coherence which match those observed in patients with DOC.^6,7^ Behaviorally, propofol anesthesia lowers both arousal and awareness, consistent with the presentation of DOC.^7^ Anesthesia has been used in several studies as a model of DOC, both in rodents and non-human primates.^8,9^ Several treatments have been explored for DOC. Amantadine moderately increases the chances that a person will emerge from a persistent DOC.^10^ Transcranial direct current stimulation and transcranial magnetic stimulation have produced inconsistent results in treating DOC.^5^ Deep brain stimulation (DBS) of the central thalamus has been shown to moderately increase metrics of arousal in human subjects with DOC, but with considerable heterogeneity in the treatment outcomes.^11^ In a study of anesthetized macaques, DBS of the CLT immediately woke them from propofol anesthesia while stimulation was on, and, in a similar study, greatly increased signs of arousal across several metrics including body movement, mouth/tongue movement, eye movement/opening and vital sign changes. ^12,13^

Recently, focused ultrasound neuromodulation (FUS) has been explored as a possible new treatment for DOC. The earliest work exploring this approach was in rodents.^8^ These experiments showed that sonication of the thalamus reduced the time to wake up from anesthesia.^8^ However, these experiments have several important limitations in their translatability to human DOC. First, the 4 × 6 mm focus of the ultrasound was large relative to the size of the rat brain (∼ 1/4 of the brain).^14^ This makes it difficult to attribute the effect of the sonication to any specific brain region. Another important limitation was the auditory confound, which, since the publication of those studies, has been shown to be a significant element in need of disentangling from the effects of FUS.^15–17^ No auditory masking or active control conditions were utilized to account for the auditory confound in these rodent studies.^8^ Studies of FUS to date in human patients with DOC have shown promising results, but have been done without sham or control groups, nor auditory masking.^18–20^ For this reason, it is not possible to confidently differentiate the effects in humans from the placebo effect or other confounds. The studies to date in humans also did not have robust patient-specific target validation, and relied on an estimated alignment between the focus of the ultrasound transducer (assumed to be unaffected by the skull) and of the location of the thalamus as determined by MRI of the patient.^18–20^

The approach taken in the following experiments addresses several of these key limitations of the previous literature. First, we used an anesthetized non-human primate model of DOC that allowed for both no-sonication and active control conditions, which allows us to disentangle any observed effects from the auditory confound. Second, we leveraged prior MR thermometry-based target validation at a nearby thalamic structure using this sonication setup to give us good confidence in FUS targeting.^21^

Safety of FUS in a model of DOC is a key consideration. Neuromodulation of the central thalamus could plausibly deepen unconsciousness and thus slow or stop breathing or the heart. While stimulation of the central thalamus with DBS has been shown to increase signatures of consciousness, the directionality of neuromodulation with FUS is still unclear, and seems both parameter and brain-region dependent, making it difficult to predict which way to expect FUS to modulate consciousness when sonicating the CLT.^22,23^ A key goal of this study was to investigate the safety of FUS of the CLT, as measured by changes in heart rate, breathing, movement and EEG.

In this study, we hypothesized the following:

- No adverse health events will occur during or after the anesthesia sessions. This will be reflected by changes in heart rate, breathing rate and blood oxygen saturation staying within clinically safe ranges, and no changes in behavior after the sessions.
- FUS of the CLT under propofol anesthesia will increase arousal. This will be reflected by increased movement, heart rate, breathing rate and increased EEG beta power.

## Materials and Methods

### Animals and Anesthesia

All in vivo procedures were approved by the University of Utah Institutional Animal Care and Use Committee (protocol number 21-12012 and 00001361) and were conducted in accordance with the Animal Welfare Regulations^24^ and the Guide for the Care and Use of Laboratory Animals.^25^ These studies were conducted within a fully AAALAC accredited facility. Two 10-year-old male Indian origin rhesus macaques (*Macaca mulatta*), subject C (13.3 kg) and B (15.2 kg) were used for this study. Both animals were chair acclimated and trained to present the lower leg for awake intravenous propofol (200 mg/20 mL injectable emulsion) (Henry Schein Inc., Melville, NY) administration through vascular access ports (VAP) attached to chronic indwelling central catheters (Cat. No. PHSPAC-3.5NC, CNC-3.5H/LSA, Access Technologies, Skokie, IL).^26^

Animals were fasted 12–16 hours prior to each experiment. Each session began with a 40 mg (4 mL) intravenous induction bolus of propofol delivered through the VAP over 30–45 seconds. Immediately following induction, intravenous propofol (subject C = 0.133 mg/kg/min, subject B = 0.139 mg/kg/min) was continued as a constant rate infusion (CRI) through the VAP for one hour to maintain a steady plane of unconsciousness. Propofol administered as a CRI was diluted 1:2 with sterile lactated Ringer’s solution (McKesson Medical Supply, Irving, TX) to increase the total intravenous fluid volume delivered per minute to avoid backflow of central venous blood into the drug line. The animal’s leg containing the VAP and propofol CRI was manually supported for the duration of the session to avoid potential damage or dislodgement in the event of movement in response to sonication. VAPs and indwelling catheters were flushed with 10 mL sterile saline, followed by 10 mL heparin (10 U/mL) (Becton Dickinson, Franklin Lakes, NJ) and locked with 0.5 mL taurolidine-citrate solution (Access Technologies, Skokie, IL) at the end of each session.

Sessions were either active (with sonication) or control (no sonication). Each sonication session was paired with a control session which was conducted at the identical propofol infusion rate. Vital parameters and clinical health were monitored by a veterinarian during and for at least 24 hours following each session.

### Transducer Setup

Ultrasonic neuromodulation was delivered using the Remus system. ^27^ This system provides a way to deliver reproducible neuromodulation of the deep brain with FUS in head-fixed rhesus macaques. Briefly, a 256-element phased array transducer is inserted into a frame that is mounted onto 4 titanium pins attached to the animal’s skull. This mounting system has been previously validated to produce reproducible targeting with this particular transducer setup.^27,28^ The transducer is coupled to the head of the animal via a cryogel in combination with standard ultrasound gel. The coupling quality is validated prior to each session using an ultrasound imaging sequence, also described previously.^27^ The ultrasound was delivered into two deep brain targets, the left and right CLT of both brain hemispheres. The experimental setup is shown in Figure 1.

**Figure 1:**
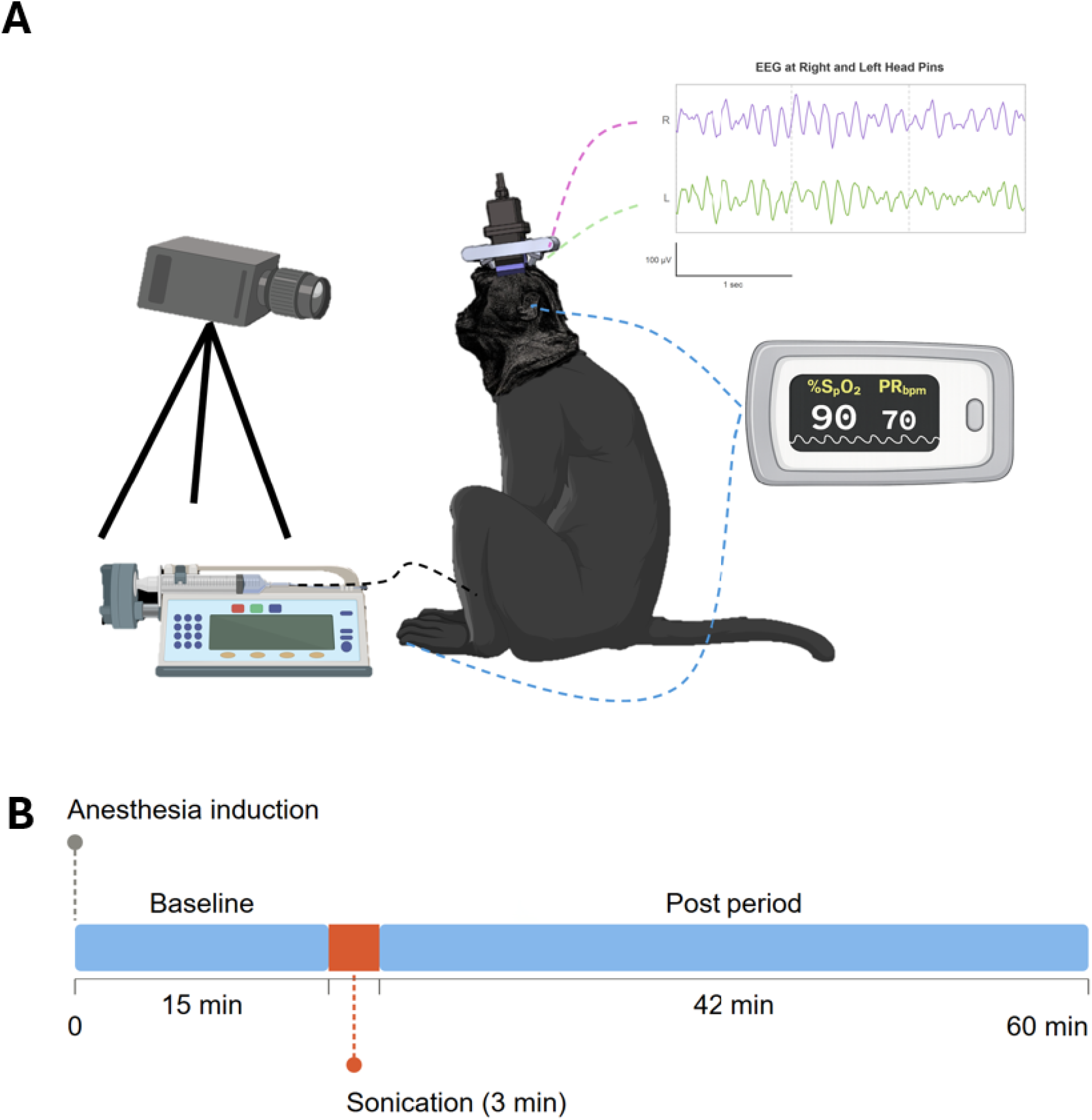
Experimental setup. (a) The Remus neuromodulation system enables repeatable sonication of deep brain circuits with focused ultrasound in head-fixed macaques, sitting upright in a primate restraint chair. ^27^ Propofol anesthesia was delivered with an infusion pump at a constant rate throughout the session. The animal was monitored via video feed, EEG, and pulse oximetry of the ear and foot. (b) Experimental protocol, repeated for every session of this experiment, with either sonication or no sonication control.

**Figure 2:**
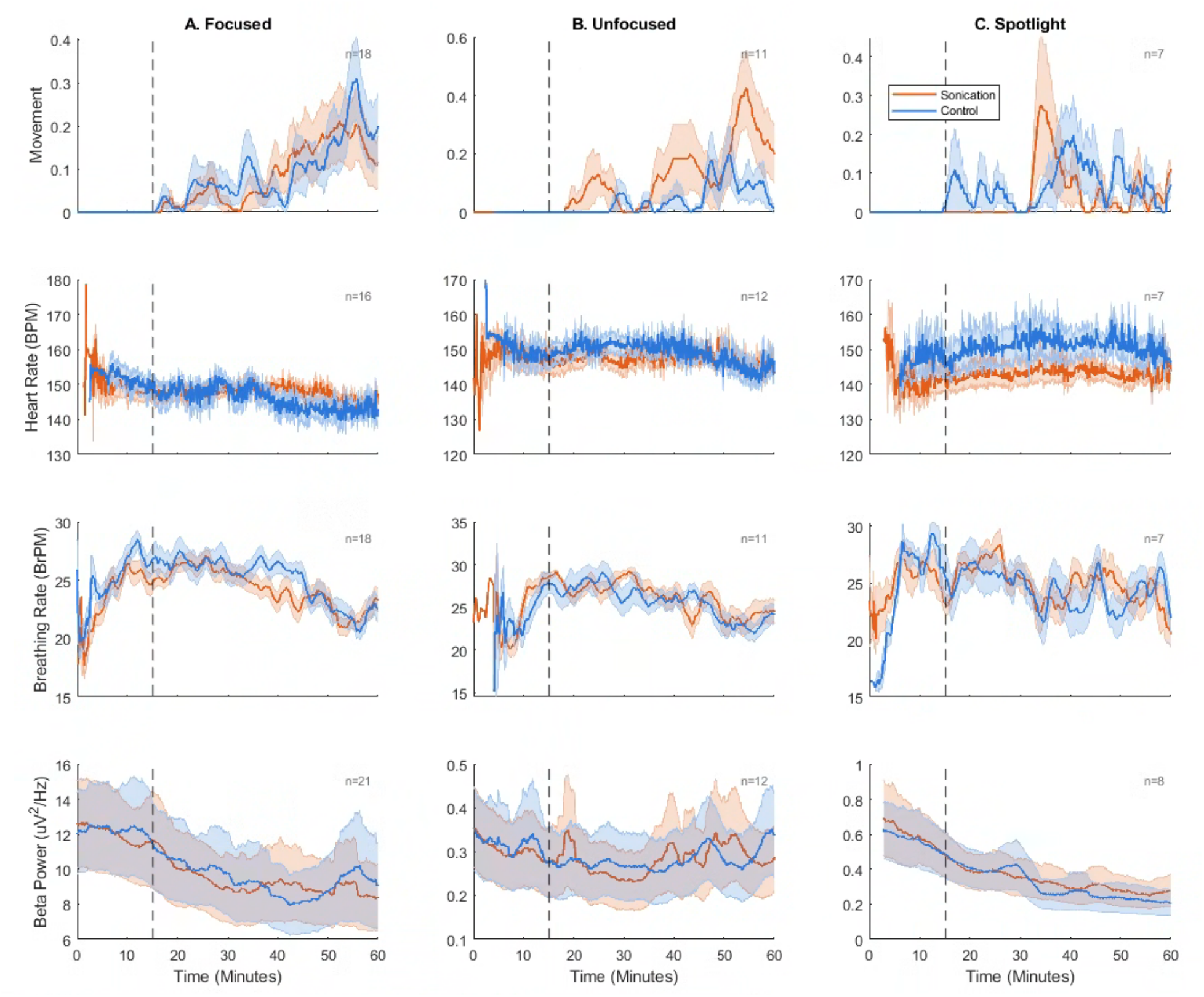
Effects of three distinct ultrasound protocols. Time-series of behavioral and physiological measures during anesthesia sessions (mean ± standard error of the mean [SEM]). Dashed vertical lines mark sonication onset. No statistically significant differences between sonication and control post-sonication were observed for any metric under any paradigm.

**Figure 3:**
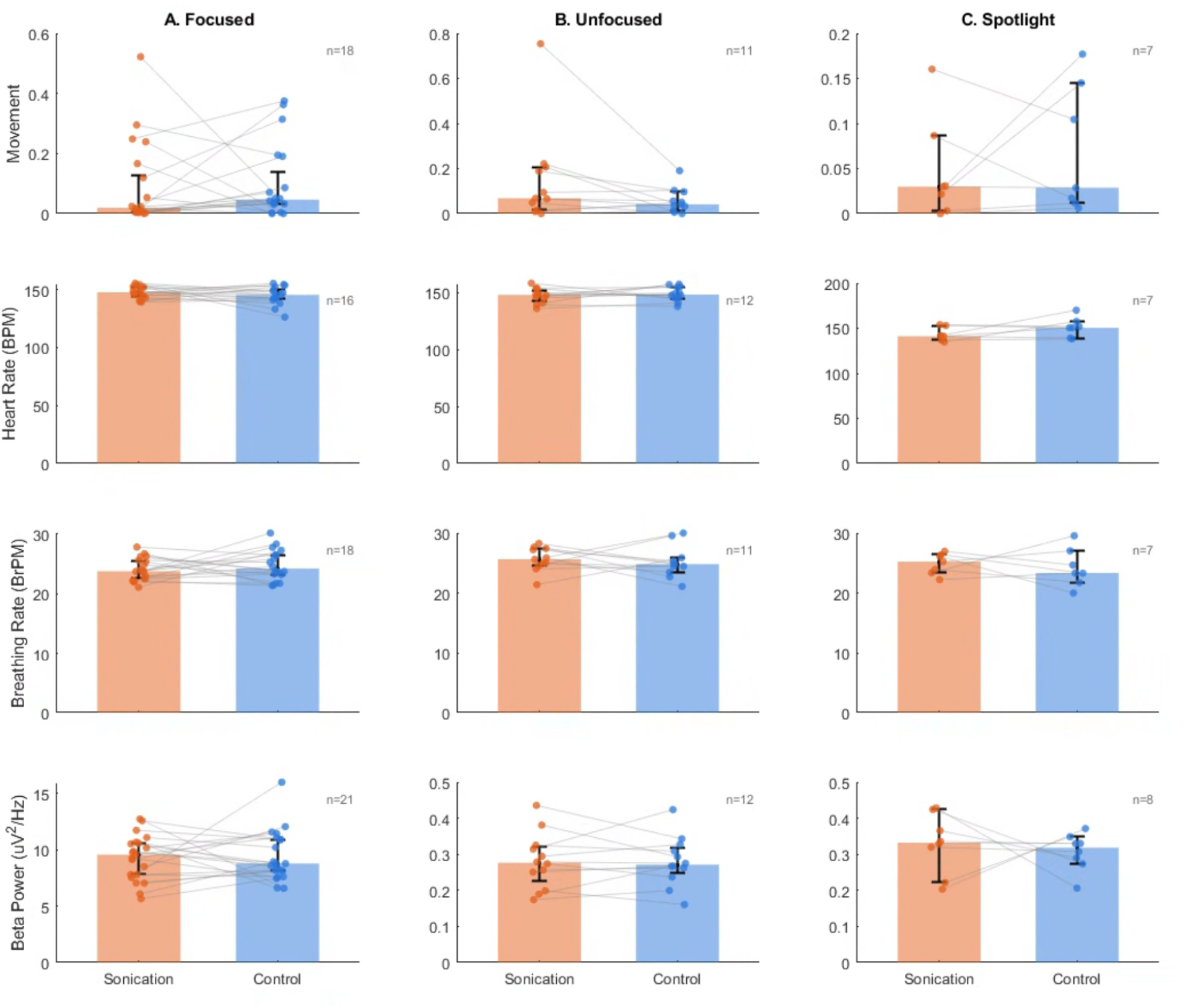
Summary of the effects. Bars show the median of each condition with bootstrap 95% confidence intervals; individual per-session post-sonication means are overlaid as scatter points with lines connecting matched pairs. Paired Wilcoxon signed-rank tests revealed no significant differences between sonication and control for any measure under any paradigm (all *p*_FDR_ *>* 0.30).

### Targeting

Targeting of the lateral geniculate nucleus (LGN) of the thalamus was validated in a previous study with this sonication setup with these animals.^27,28^ From the LGN, we measured the distance from the center of the LGN to the CLT using a brain atlas, and steered the ultrasound beam the commensurate three-dimensional distance to target the CLT. Direct target validation was avoided here because MR thermometry (the method our group has used for target validation in other experiments) involves depositing large amounts of energy into the brain to heat the tissue, which would present a substantial safety risk in a brain region controlling consciousness, such as the CLT. We have validated via thermometry in a previous experiment that steering from the LGN to the amygdala was done with ± 1 mm error, giving us good confidence in the steering accuracy to the CLT, a much closer relative target.

### Ultrasound Neuromodulation Parameters

Three sonication paradigms were tested: focused, unfocused, and spotlighting. For all three paradigms, sonications used a 50% duty cycle, 50 Hz pulse repetition frequency (PRF), 1 MPa estimated in-situ peak rarefactional pressure, 480 kHz carrier frequency, and 300 ms pulse duration. In-situ pressure was estimated according to previously described methods.^27^ Briefly, in-situ intensity was estimated to be 21% of the free field intensity, based on MR thermometry. The sonication parameters were chosen as an adaptation of the parameters that we have previously used to see behavioral effects in our non-human primates, combined with a 50 Hz PRF to replicate the pulse frequency used in the studies using DBS to awaken non-human primates from anesthesia.^12,28^ We expected that matching the PRF to the frequency of electrical stimulation in previous studies would produce behavioral effects in the same direction.

In the focused condition, bilateral CLTs were sonicated in alternating 3-minute blocks, with a 3 s inter-stimulus interval between successive sonications of each CLT. Each CLT was sonicated 40 times per block. This resulted in an *I*_SPTA_ of 1.6 W/cm^2^ and an *I*_SPPA_ of 32.3 W/cm^2^, with a corresponding mechanical index (MI) of 1.44. This MI is within the ITRUSST consensus limit of 1.9.^29^ The *I*_SPPA_ was also within the FDA 510(k) Track limits for diagnostic ultrasound, while the *I*_SPTA_ limit was exceeded.^30^ It has been well established in previous studies in both macaques and humans that this *I*_SPTA_ limit can be safely exceeded for ultrasound neuromodulation, and an *I*_SPTA_ of up to 25.8 W/cm^2^ has been used in macaques without tissue damage. ^31^

In the unfocused condition, the phases of the 256 array elements were randomized, spatially dispersing the acoustic energy across the brain rather than focusing it at the CLT. The total acoustic energy delivered and all pulse parameters were equivalent to the focused condition. This condition served as a control for non-focal effects of ultrasound delivery, including potential auditory confounds.

In the spotlighting condition, the ultrasound beam was sequentially steered to four positions located 2 mm diagonally from the central CLT target, alternating bilaterally as in the other conditions. Successive positions were sonicated 30 seconds apart, with each position receiving 10 sonications. All pulse parameters were identical to the focused condition, resulting in the same *I*_SPPA_ (32.3 W/cm^2^) and an *I*_SPTA_ of 0.4 W/cm^2^ at each position. This condition tested whether sonicating regions adjacent to the CLT would stimulate nearby tracts or nuclei that could produce an acute effect on arousal, and to account for minor error in targeting accuracy.

### Session Structure

Each animal was anesthetized in 1-hour sessions once per week, with at least three days between sessions. 15 minutes after the beginning of the session, the animal either received sonication or no sonication for the session. The animal was then monitored for the next 45 minutes of the anesthesia session.

### Monitoring of Consciousness

Consciousness was monitored throughout the duration of each anesthesia session via EEG, pulse oximetry (heart rate and oxygen saturation), and video to record movements and breathing rate. Significant changes in blood oxygen saturation were documented manually during the session, with the time of any significant changes noted by the experimenter.

Pulse oximetry was recorded from sensors on the ear and foot. Signals were bandpass filtered at 1-5 Hz for cardiac peak detection, and heart rate was estimated from inter-peak intervals within a sliding 5-second window. When both sensors were available, the channel with the more stable heart rate estimate was automatically selected in 75-second blocks. If one channel showed near-zero variability (range < 0.5 BPM), indicating sensor dropout, the other channel was substituted for that segment. Sessions with greater than 50% missing data were excluded.

EEG beta power was calculated (13–30 Hz) using 0.5 s windows with 50% overlap from the two rear headpins of the animal, roughly corresponding to P3/P4 positions in the 10/20 EEG system. The two frontal pins served as ground and reference. An Intan RHS2000 EEG acquisition system was used, which sampled the signals at 20 kHz and low-pass filtered the signal at 7.5 kHz. The average impedance of the EEG recording was 0.30 kΩ. Samples with absolute activity greater than 150 *µ*V were considered spurious and were removed from the analysis. A notch filter was applied at 60 Hz. The EEG channel with the lowest standard deviation for each session was selected. This was done because during movement, occasionally an electrode was shaken loose, resulting in extreme persistent high noise on that channel.

Video monitoring of the movement of the animal was analyzed in Python using the OpenCV library. OpenCV is a computer vision library designed to automatically analyze and quantify changes in video footage.^32^ OpenCV-based pipelines are capable of achieving human-level accuracy in quantifying movement in animal behavioral tasks, reporting a strong correlation of up to *R*^2^ = 0.98 when compared to manual scoring.^33^ Movement was quantified by manually selecting the animal’s body as a region of interest, and then taking pixel differences via subtraction frame-to-frame of the video. A 5-minute period before sonication was used as the baseline with which to compare movement. Periods in which quantified movement had a range that was 2 standard deviations above the baseline period were classified as movement. Breathing rate was extracted from the general movement data using a 0.167–0.667 Hz (10–40 breaths per minute) bandpass filter, which represents the range of breathing rate for lightly anesthetized macaques.^34^

### Statistical Analysis

Statistical analysis was performed with a custom Matlab script (Matlab R2024a). For each physiological measure (EEG beta power, heart rate, movement, and breathing rate), session-level summary statistics were computed as the mean value over the post-sonication period within the session. Sonication and control sessions were paired within animal and infusion rate using greedy nearest-date matching. Differences between sonication and control conditions were assessed using two-sided paired Wilcoxon signed-rank tests applied to per-pair (sonication minus control) differences, for each measure within each sonication paradigm (focused, unfocused, and spotlighting). Effect sizes were quantified as matched-pairs rank-biserial correlation, with 95% confidence intervals obtained from 5,000 percentile bootstrap resamples. Raw *p*-values were corrected for multiple comparisons across the three paradigms within each measure using the Benjamini–Hochberg false discovery rate procedure. Each measure was treated as a separate test family. To assess consistency across subjects, paired Wilcoxon signed-rank tests were also performed separately within each animal, with false discovery rate correction applied across paradigms within each animal/measure combination.

### Omitted Data

Several sessions were omitted from specific analyses due to partial or complete data loss during the session. For heart rate, sessions with *>*50% missing pulse oximetry data were excluded: 3 sonication sessions and 1 control session in the focused condition, and 1 control session in the spotlighting condition. No heart rate sessions were excluded in the unfocused condition. For movement and breathing rate, several sessions lacked video recordings due to equipment failures and were excluded from those analyses. These data losses were metric-specific; a session excluded from heart rate analysis due to pulse oximetry failure could still contribute valid EEG, movement, and breathing data, and vice versa. As a result, the number of valid pairs varied by metric within each paradigm. Additionally, 3 control sessions in the focused paradigm had no matching sonication session at the same animal and infusion rate and were thus excluded from pairing. The spotlighting paradigm contains sessions from subject B only; no spotlighting sessions were conducted on subject C.

## Results

### Post Sonication Results

Sonication and control sessions were paired by animal and infusion rate, yielding 21 pairs in the focused paradigm (subject C 13, subject B 8), 12 pairs in the unfocused paradigm (6 per animal), and 8 pairs in the spotlighting paradigm (subject B only). Due to metric-specific data losses described above, heart rate analyses used 16, 12, and 7 pairs, and movement and breathing rate analyses used 18, 11, and 7 pairs for the focused, unfocused, and spotlighting paradigms, respectively. Across all three paradigms, paired Wilcoxon signed-rank tests revealed no significant differences between sonication and control conditions for any of the four physiological measures (all *p*_FDR_ *>* 0.30). Matched-pairs rank-biserial effect sizes were small and inconsistent in direction across paradigms and measures, ranging from −0.36 to +0.58 with 95% bootstrap confidence intervals that crossed zero in every case. The largest observed effect was an increase in body movement under the unfocused paradigm (*r*_rb_ = +0.58, 95% CI [− 0.09, +0.94]; *p*_raw_ = 0.10, *p*_FDR_ = 0.30), though this did not reach statistical significance. Per-animal analyses confirmed the absence of systematic effects, with neither subject B nor subject C showing significant differences individually in any paradigm after false discovery rate correction (all *p*_FDR_ *>* 0.32).

### Safety Results

The animals remained physiologically stable and free of clinical health concerns during the experimental sessions and subsequent recovery periods. Body weight was not significantly altered in either animal while on study. During the experiments, heart rate never dropped below normal ranges, breathing remained constant and blood oxygen remained clinically stable throughout all sessions.

## Discussion

In this study we hypothesized that we would find no adverse effects of FUS of the CLT and that FUS of the CLT would increase signatures of arousal. The results of this study showed no safety concerns with FUS of the CLT under the studied sonication protocol, which could have plausibly either increased or decreased arousal. Decreased arousal could dangerously suppress heart rate and breathing. ^7,13^ No results indicated safety challenges, including no notable drops in heart rate, breathing, or blood oxygen saturation during or after the sonication. We also found no significant modulation of arousal across any metric after FUS of the CLT. No significant changes in body movement, heart rate, breathing rate or EEG beta power were detected across either animal under any sonication paradigm.

These results indicate that FUS of CLT may have limited effectiveness for increasing arousal for those suffering from DOC, at least with the tested sonication paradigms. The largest observed effect was an increase in movement under the unfocused paradigm (*r*_rb_ = +0.58, 95% CI [−0.09, +0.94]), which had a moderate effect size, but did not reach statistical significance. This paradigm distributed equivalent acoustic energy across the thalamus, rather than focusing it at the CLT, and may indicate that distributed thalamic sonication could induce broader network modulation effects relevant for increasing consciousness, which is itself a distributed brain-wide phenomenon^6,7^.

This study extends prior work by investigating FUS of the CLT in a large-animal model of DOC. This study includes several improvements over previous experiments. Specifically, compared with the rodent literature sonicating the thalamus (but which sonicated approximately 1/4 of the brain),^8,14^ our study sonicated a much more focused brain region, relative to the size of the brain of the animal. Second, we included an unfocused ultrasound condition, which, if there was an effect due to auditory confounds, would have produced an effect on its own, separate from the effects of the focused ultrasound.^17^ In sum, relative to the previous rodent literature, we investigated sonication of a more focal brain region and addressed the auditory confound.

Previous proof of concept studies investigating FUS of the thalamus of human subjects with DOC were crucially missing control groups or sham conditions.^18–20^ In these studies the subjects showed greater recovery from DOC than was expected had there been no intervention. Without a control or sham group, it is impossible to disentangle these effects from possible confounds, such as increased recovery due to touching the patient, hearing voices of the researchers, etc. These studies also had limited target validation. In these studies the focal distance of the transducer was used to estimate the approximate focus of the ultrasound in the brain as the transducer was placed on the skull.^18–20^ Our study added to this body of work sonication of the thalamus in a model of DOC, with robust control/sham conditions, and a greater degree of target validation (validation done in the LGN of the thalamus via MR thermometry and electronically steered using phase interference to the CLT, also in the thalamus).

This work includes several key limitations. First, the targeting of the CLT was done by first validating targeting at the LGN in the thalamus from previous studies, and then steering electronically via phase interference to the CLT of the thalamus. This steering has been shown in a previous study to be relatively robust, where we steered to the amygdala, and the distance input into the software steering of the transducer matched ±1 mm to the center of the new brain region as validated by MR thermometry.^21^ There was also uncertainty in the delivered ultrasound intensity. Our estimate, based on MR thermometry, relies on assumptions about the thermal properties in the brain, and both spatial and temporal averaging inherent to MR thermometry.^27^ Another limitation of this work is that it only included two animals, which limits both the statistical power and generalizability of the results to other animals and species. This small number of animals was for practical reasons, including the number of available animals. To preserve the health of the animals and allow for adequate recovery, sessions were performed at 1 session per week. This limited the number of experimental protocols and sonication parameter sets that could be tested.

The results of this study contrast to those observed with DBS of the CLT in macaques. DBS showed dramatic and reliable arousal responses that corresponded to near-complete behavioral separation from the anesthetized state.^12,13^ These experiments showed the animals opening their eyes, moving their limbs, and showing waking vital sign changes. ^12,13^ In contrast, our results using FUS at the same target showed no statistically significant differences between sonication and no sonication. Matched-pairs rank-biserial effect sizes for the focused paradigm were also small in magnitude (|*r*_rb_| ≤ 0.22 across all four measures), with 95% bootstrap confidence intervals of approximately ±0.5 that included zero. These intervals are consistent with either no effect or small true effects. A larger sample size would be required to confidently rule out small effects of FUS at the CLT under the tested sonication parameters. Regardless, the effects observed here do not approach those observed with DBS of the CLT in macaques under propofol anesthesia.

Future work exploring FUS as a treatment of DOC should investigate the role of sonication spatial distribution. The largest (though not statistically significant) effect in this study occurred under the unfocused paradigm, which may suggest that engaging a broader thalamic network could be more effective at promoting wakefulness than focal targeting of individual nuclei with FUS. Higher pressures and sonication durations than those tested here (which were selected to be relatively conservative) could also increase neuromodulatory effectiveness of focused or unfocused paradigms. Future studies should exercise caution to include both robust control conditions and robust target validation, which were missing from prior work.

## Acknowledgments

None.

## Data Availability

Supporting data will be made available on request.

